# Relationships between the Bone Expression of Alzheimer Disease-Related Genes, Bone Remodelling Genes and Cortical Bone Structure in Neck of Femur Fracture

**DOI:** 10.1101/2020.11.02.365866

**Authors:** Catherine J.M. Stapledon, Roumen Stamenkov, Roberto Cappai, Jillian M. Clark, Alice Bourke, L. Bogdan Solomon, Gerald J. Atkins

## Abstract

Neck of femur (NOF) fracture is a prevalent fracture type amongst the ageing and osteoporotic populations, commonly requiring total hip replacement (THR) surgery. Increased fracture risk has also been associated with Alzheimer disease (AD) in the aged. Here, we sought to identify possible relationships between the pathologies of osteoporosis and dementia by analysing bone expression of neurotropic or dementia-related genes in patients undergoing THR surgery for NOF fracture. Femoral bone samples from 66 NOF patients were examined for expression of the neurotropic genes amyloid precursor protein (*APP*), APP-like protein-2 (*APLP2*), Beta Secretase Cleaving Enzyme-1 (*BACE1*) and nerve growth factor (NGF). Relationships were examined between the expression of these and of bone regulatory genes, systemic factors and bone structural parameters ascertained from plain radiographs. We found strong relative levels of expression and positive correlations between *APP, APLP2, BACE1* and *NGF* levels in NOF bone. Significant correlations were found between *APP, APLP2, BACE1* mRNA levels and bone remodelling genes *TRAP, RANKL*, and the *RANKL:OPG* mRNA ratio, indicative of potential functional relationships at the time of fracture. Analysis of the whole cohort, as well as non-dementia and dementia sub-groups, revealed structural relationships between *APP* and *APLP2* mRNA expression and lateral femoral cortical thickness. These findings suggest that osteoporosis and AD may share common molecular pathways of disease progression, perhaps explaining the common risk factors associated with these diseases. The observation of a potential pathologic role for AD-related genes in bone may also provide alternative treatment strategies for osteoporosis and fracture prevention.

## Introduction

In Australia, the proportion of individuals aged 65 and over has increased from 12.3% to 15.9% in a twenty year period [1]. This has resulted in a parallel rise in the rate of development of age-related pathologies. Neck of femur (NOF) fracture is a painful and debilitating fragility-related fracture, that negatively impacts the lives of over 19,000 Australians every year [2]. The risk of developing fragility fractures, such as NOF, increases substantially with increasing age [3]. Other risk factors associated with the development of a NOF fracture include gender, with females being at greater risk than males, low bone mineral density (BMD) attributable to osteopenia or osteoporosis [4, 5] and prior fracture [6]. Osteoporosis is one of the most common musculoskeletal diseases associated with increased fracture risk. The epidemiology of dementia mirrors that of osteoporosis. Dementia is the second leading cause of death and disability in Australia [7, 8] and shares many of the same risk factors as fracture and osteoporosis. The most commonly occurring form of dementia in Australia is Alzheimer’s disease (AD), which accounts for up to 70% of all dementia diagnoses. There are currently no effective therapeutics for the prevention or treatment of AD.

Previous studies have sought to determine links between increased fracture risk and the development of AD and other forms of dementia [9-15]. A number of risk factors have been identified through observational cohort studies linking these pathologies, including age, gender, vitamin D status, bone mineral density (BMD) and comorbidities, such as chronic kidney disease [16]. Data from the Australian New Zealand Hip Fracture Registry indicate that while 59% of hip fracture patients reported no problem with cognition prior to hospital admission, 41% of such patients had either not been assessed for cognition prior or were suffering from a cognition-related illness [17]. Recent studies have reported that patients with a low BMD have a higher incidence of AD and subsequent risk of fracture, however underlying mechanisms linking these major health issues are yet to be determined [18-22].

Hip fractures associated with frailty and OP usually occur at the femoral neck (56%), pertrochanteric region (38%) or subtrochanteric region (5.8%). They are often a result of a fall or minimal trauma, associated with poor bone quality and strength. The cellular mechanisms underlying fragility fracture are associated in part with dysregulated bone remodelling leading to net bone loss. The RANKL/RANK/OPG signalling pathway plays a prominent role in this process, controlling the formation and activity of the key bone resorbing effector cell in the bone remodelling process, the osteoclast [23]. However, low bone formation rates and poor bone quality at key skeletal sites, as well as muscle quality and mass are other important determinants of bone strength and resistance to low trauma fracture. Osteocytes have emerged as an important controlling cell type in various aspects of bone remodelling, as well as in a number of systemic roles [24, 25]. The presence of an intact osteocytic network is also important in the maintenance of bone integrity and this degrades with ageing and the onset of osteoporosis [26-28]. Osteocyte-derived factors, such as RANKL and sclerostin, encoded by the *SOST* gene, have emerged as important therapeutic targets in the treatment of osteoporosis [24]. Osteocyte control of RANKL-dependent osteoclastogenesis has been shown in response to mechanical [29, 30], as well as metabolic and dietary signals [31, 32]. In adult human bone, osteocytes are a prominent source of RANKL [33]. We have shown that pathological stimuli, such as orthopaedic implant derived wear particles also likely exert an effect through this pathway [34]. Human osteocytes have also been reported to express neurotropic factors, including nerve growth factor (NGF) [35] and the amyloid precursor protein (APP) [36]. The study by Li and colleagues [36] also reported osteocyte expression of the neurotoxic peptide Aβ_1-42_, derived from post-translational processing of APP by the beta-secretase BACE1, and which is a known contributor to AD pathogenesis.

In this study, we sought to uncover links connecting the pathologies of osteoporosis and dementia by investigating the bone expression of AD-related and neurotropic genes in bone taken from patients undergoing surgery for NOF fracture, and relating this to bone turnover-related gene expression, as well as bone structure. We present evidence for novel relationships between neuropathic gene expression, bone remodelling and bone structure, potentially contributing to fragility fracture and reflecting cognition in this cohort of patients.

## Materials and Methods

### Sample Collection

Intertrochanteric trabecular bone biopsies, plasma and pre-surgical radiographs were collected from patients undergoing hip arthroplasty for a fracture NOF at the Royal Adelaide Hospital (RAH) Adelaide, South Australia, between July 2018 and July 2019. During preoperative preparation, patients or next of kin, depending on cognitive state, were asked to participate in this study, and informed written consent was obtained. Date of surgery, patient age, comorbidities, creatinine levels, vitamin D status and medication history were collected from patient case notes. The study was approved by the Human Research Ethics Committee of the RAH (Protocol no 130114; HREC/13/RAH/33).

### Cohort Demographics and Comorbidities

Participants were screened upon hospital admission prior to surgery and daily thereafter using the Confusion Assessment Method (CAM) in association with the Abbreviated Mental Test Score (AMTS) [37]. Perioperatively, participants were assessed for cognitive impairment using the Mini-Mental State Examination (MMSE) [38]. In some cases, participants had a diagnosis of AD or dementia prior to hospital admission. Participants were excluded if their bone samples yielded poor RNA quality or amount (260:280nm absorbance ratio < 1.7; yield < 100 ng/µl). Thus, of the 93 consented participants who provided bone biopsies and plasma samples, 66 were included for further analysis, including 53 females (80.3 %) and 13 males (19.7 %). Comorbidity data was obtained from the SA Health OACIS Medical Record Database from the RAH. Demographic and comorbidity data are shown in **Table 1**.

**Table 1:** Whole Cohort Correlation Analysis of Gene Expression with Structural Parameters. Relative gene expression was generated by Real-time RT-PCR normalised to 18S rRNA levels. Structural parameters measured from plain radiographs: Lateral cortex (LC); Medial cortex (MC); Femoral canal (FC); Proximal femur outer (PFO). Significant correlations and their respective *p* values are bolded.

| *Gene* | *LC (1)* |  | *MC (2)* |  | *FC (3)* |  | *PFO (4)* |  |
| --- | --- | --- | --- | --- | --- | --- | --- | --- |
|  | *r* | *p* | *r* | *p* | *r* | *p* | *r* | *p* |
| *APP* | **-0.329** | **0.007** | -0.068 | 0.585 | -0.021 | 0.866 | -0.152 | 0.224 |
| *APLP2* | **-0.354** | **0.004** | -0.116 | 0.359 | -0.007 | 0.958 | -0.182 | 0.147 |
| *BACE1* | -0.196 | 0.118 | -0.090 | 0.478 | -0.032 | 0.798 | -0.137 | 0.276 |
| *CYP27B1* | 0.196 | 0.120 | -0.007 | 0.955 | 0.140 | 0.270 | 0.175 | 0.166 |
| *DMP1* | **-0.253** | **0.042** | 0.053 | 0.674 | -0.012 | 0.923 | -0.103 | 0.415 |
| *NGF* | -0.196 | 0.276 | -0.157 | 0.384 | 0.045 | 0.804 | -0.104 | 0.565 |
| *OCN* | -0.032 | 0.802 | 0.011 | 0.930 | 0.014 | 0.910 | 0.020 | 0.872 |
| *OPG* | **-0.313** | **0.014** | 0.030 | 0.817 | 0.034 | 0.792 | -0.070 | 0.590 |
| *RANKL* | **-0.385** | **0.002** | -0.096 | 0.467 | -0.078 | 0.553 | **-0.294** | **0.023** |
| *RANKL:OPG* | -0.142 | 0.279 | -0.114 | 0.386 | -0.217 | 0.095 | **-0.358** | **0.005** |
| *SOST* | 0.120 | 0.544 | 0.140 | 0.477 | 0.243 | 0.213 | 0.317 | 0.100 |
| *TRAP* | **-0.316** | **0.013** | -0.069 | 0.596 | -0.127 | 0.329 | -0.247 | 0.054 |

### Processing of Bone Biopsies

During surgery tube saw cancellous bone biopsies were taken from the intertrochanteric region before the femoral canal was prepared for receiving the femoral stem. Samples were submerged in saline in a sterile container and collected from theatre within 48 hours of surgery. All samples were processed in a sterile biosafety cabinet. Each sample was cut into approximately 2 cm^3^ cubes and washed 3-4 times with sterile 1 x phosphate buffered saline (PBS) to remove loosely adherent cells and debris. Samples were then stored at -80°C for up to two weeks before processing for RNA. Samples for histology were stored in Osteosoft® solution (Merck KGaA; Darmstadt, Germany), for one week and processed for histological analysis.

### RNA Extraction

Bone biopsies stored at -80°C between 1 and 7 days were placed into liquid nitrogen contained in a ceramic mortar sprayed with RNaseZap^®^ (ThermoFisher Scientfic, MA, USA). Samples were pulverised using an RNaseZap-treated pestle. Bone remnants were transferred into a 1.5 ml RNAse-free reaction tube (Eppendorf AG, Hamburg) containing 1 ml of TRIZOL reagent (Life Technologies, NY, USA). RNA extraction was conducted as per the manufacturer’s instructions [39]. Complementary DNA (cDNA) synthesis was performed using iScript™ RT kit (Bio-Rad, Hercules, CA, USA), as per the manufacturer’s instructions.

### Real-Time RT-PCR

Real-time RT PCR was conducted for amyloid precursor protein (*APP*), amyloid precursor-like protein-2 (*APLP2*), beta-secretase cleaving enzyme-1 (*BACE1*), 25-hydroxyvitamin D 1∝-hydroxylase (*CYP27B1)*, dentin matrix acid phosphoprotein-1 (*DMP1*), nerve growth factor (*NGF)*, osteocalcin (*OCN*), osteoprotegerin (*OPG)*, receptor activator of nuclear factor kappa-B ligand (*RANKL*), sclerostin (*SOST*) and tartrate resistant acid phosphatase (*TRAP*). Oligonucleotide primers were designed to be human mRNA-specific. Primer sequences are shown in **Supplementary Table 1**. Cycle threshold (CT) values were normalised to those of the *18S* housekeeping gene using the delta-CT method (2^-(CT1-CT2)) [40].

### Radiographic Analysis

Calibrated Anterior-Posterior (AP) plain radiographs of the contralateral femur were used in PACS software to measure medial (M) and lateral (L) cortical thicknesses, and M-L medullary width and femoral width using CARESTREAM software version 5.**7 (**Windsor, CO, USA).

### Statistical Analyses

All statistical testing was conducted using the GraphPad Prism Analysis Program v.7.02 (GraphPad Prism, La Jolla, CA, USA). Gene expression levels were analysed using a non-parametric, One-way Analysis of Variance (ANOVA) test, correcting for multiple comparisons. Relationships between the expression of various genes were tested using Spearman’s correlation analysis. A *p* value < 0.05 was considered significant for all analyses.

## Results

### Characteristics of participants

The mean age of the 66 patients investigated was 81.9 ± 9.15 years (range 58 – 96). Serum 25(OH)vitamin D (Vitamin D) levels ranged from 11 – 154 nM. Serum creatinine levels were widely variable, ranging from 40 – 303 pM. The most common pre-fracture comorbidities within the study population were hypertension (65.15 %), hypercholesterolemia (30.30 %), osteoarthritis (25.76 %), type II Diabetes Mellitus (25.76 %), atrial fibrillation (24.24 %), gout (24.24 %), and osteoporosis (18.18 %) (**Supplementary Table 2**). Including those identified in-hospital, 13 (19.69 %) participants had a diagnosis of dementia. In this study, there were a total of 5 in-hospital deaths (7.6%), 2 males (15.4% of the male group) and 3 females (5.7% of the female group).

### Gene expression analysis of the NOF cohort

In order to investigate potential functional relationships between genes with known roles in neuronal function or neurodegenerative disease and those with known roles in bone remodelling, real-time RT PCR analysis was performed. Initial analysis revealed high mRNA expression levels of *APP, APLP2* and of the marker of osteoclast activity, *TRAP* (**Fig. 1a**). In comparison to *APP* and *APLP2, BACE1* levels were lower although present in all bone samples. A strong correlation between *APP* and *APLP2* expression was observed (r = 0.88; **Fig. 1b**). *APP* mRNA expression also correlated with those of *BACE1* (*p* < 0.0001), *DMP1* (*p* = 0.0004), *NGF* (*p* = 0.014), *OPG* (*p* = 0.0005), *RANKL* (*p* < 0.0001), the *RANKL:OPG* mrna ratio (*p* = 0.021) and *TRAP* (*p* < 0.0001) (**Fig. 1b**). *APLP2* mRNA expression also correlated with that of *BACE1* (*p* = 0.003), however the relationship was not as strong as *APP* and *BACE1. APLP2* correlated with the same genes as *APP* (*DMP1 p* = 0.033; *NGF p = 0.0203; OPG p = 0.027; RANKL p* < 0.0001; *RANKL:OPG* ratio *p* = 0.0009 and *TRAP p* < 0.0001). There were observable correlations between *BACE1* and *DMP1* (*p* = 0.0003), *NGF* (*p* = 0.017), *OPG* (*p* < 0.0001), *RANKL* (*p* = 0.0003) and *TRAP* (*p* < 0.0001). *APLP2* was positively correlated with the same set of genes. *BACE1* was the only gene to reveal a positive correlation with age, however it did not correlate with the *RANKL:OPG* mRNA ratio, an indicator of active bone resorption (**Fig. 1b**). There was a strong negative correlation between the *RANKL:OPG* mRNA ratio and *SOST* mRNA expression (*r* = -0.659, *p* < 0.0001).

**Figure 1:**
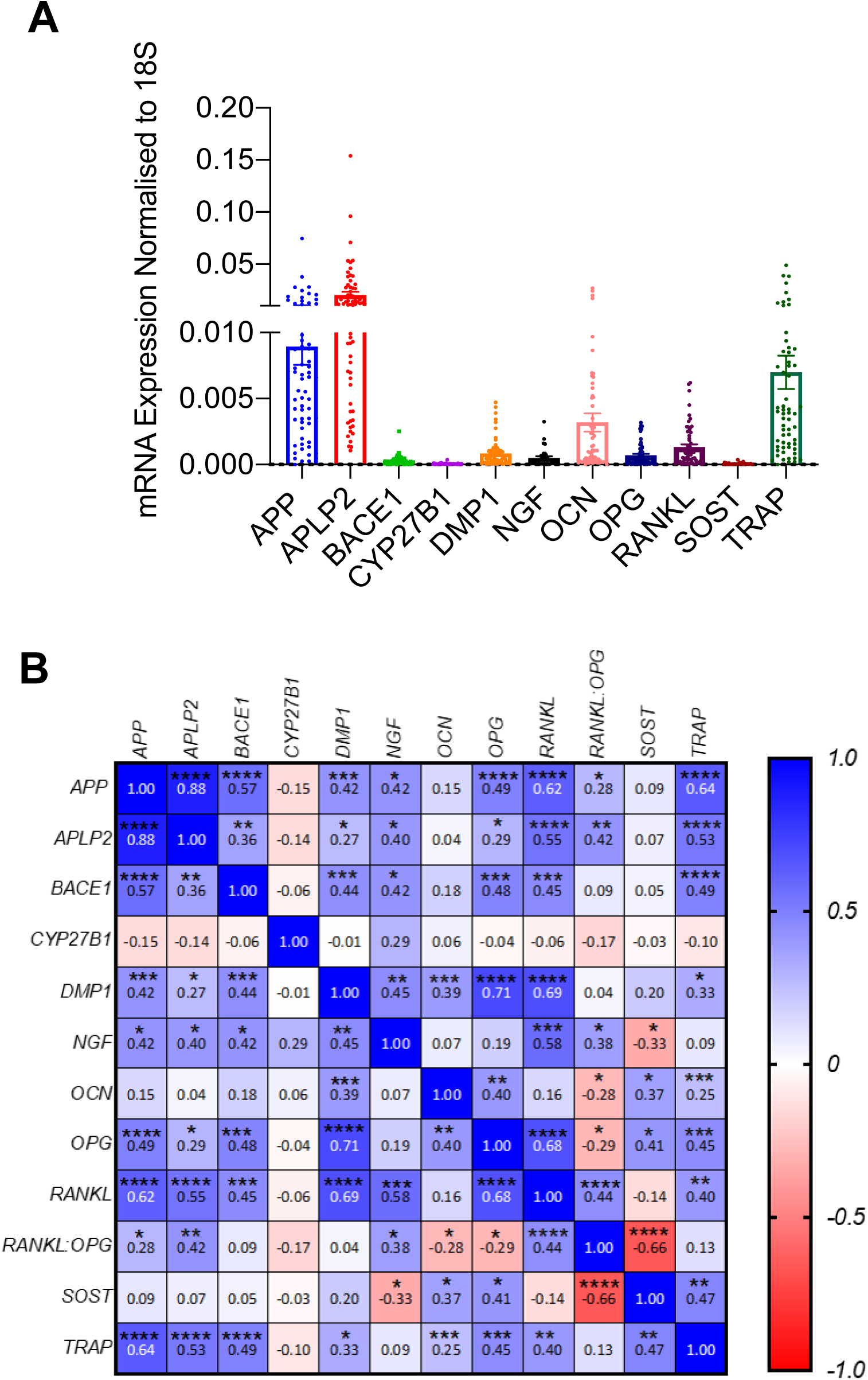
Gene expression and correlation analysis for the whole NOF cohort. A. Gene expression of each gene of interest normalised to the 18S housekeeping gene. B. A heat map representation of the relationships between each gene of interest using the Spearman’s correlation co-efficient. *r* values are reported. Relationship direction and strength are indicated by the legend: 0 – 1.0 indicates positive relationship between genes and - 1.0 – 0 indicates a negative relationship between genes. The significance of *r* values is indicated by*\*p* < 0.05, \*\**p* < 0.01 and \*\*\**p* < 0.001.

### Relationships between bone gene expression and systemic markers

Correlation analysis was performed in order to determine whether relationships existed between routinely assessed serum vitamin D and serum creatinine levels and bone cell gene expression. Whole cohort analysis did not reveal statistically significant relationships between any of the genes analysed and serum vitamin D (**Supplementary Table 3**). Serum creatinine levels, an indicator of normal kidney function [41, 42], were also tested. Interestingly, a significant positive correlation was observed between serum creatinine levels and *OPG* mRNA expression (*r* = 0.346; *p* = 0.006) (**Supplementary Table 3**). However, no other relationships between serum vitamin D or creatinine with bone gene expression were found.

### Whole cohort relationships between gene expression and bone structure

As most NOF patients did not have bone densitometry performed prior to their fracture, BMD values were not available. However, physical measurements of the proximal femur from plain radiograph images were analysed for: 1 – lateral cortical thickness, 2 – medial cortical thickness, 3 – medullary width and 4 – femoral width (**Fig. 2a - e**).

**Figure 2:**
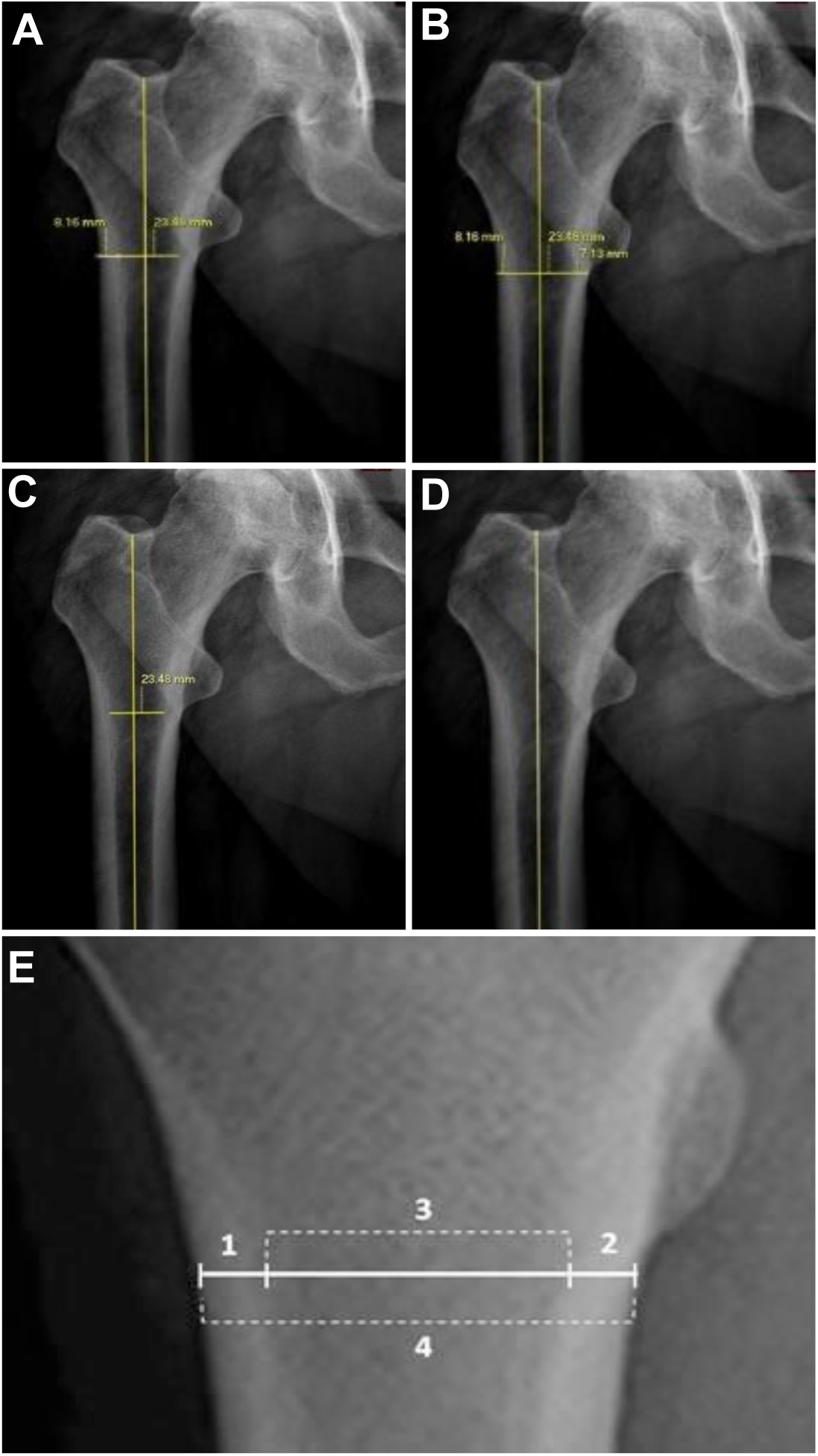
Examples of AP plain radiographic images of the hip joint with manual linear measurements applied conducted using the CARESTREAM program. (A) Measurement of the lateral cortex. (B) Measurement of the medial cortex. (C) Measurement of the femoral canal *measured from the most distal point of the lesser trochanter perpendicular to femoral midline. (D) Femoral midline measurement. (E) Measurements of the femoral neck used for analysis: 1 – Lateral cortical thickness (µm); 2 – Medial cortical thickness (µm); 3 – Medullary width (mm); 4 – Femoral width (mm).

Spearman’s *r* values were calculated between expression of genes of interest and femoral structural measures. Analysis of markers of bone remodeling revealed significant correlations between lateral cortical thickness of the femur and *RANKL, TRAP, DMP1* and *OPG* (**Table 1**). The strongest negative correlation was observed between femoral cortical thickness and *RANKL*, followed by *TRAP, OPG* and *DMP1* (**Table 1, Fig. 3**). These findings served to validate the approach taken and imply that femoral cortical thickness is causally related to the expression of these genes. There were no correlations between any of the structural parameters and *BACE1* (**Table 1**). Interestingly, negative correlations were observed between *APP* and *APLP2* mRNA expression and lateral cortical thickness (**Table 1, Fig. 3**). This suggests that expression of these genes is associated with loss of cortical bone thickness through influences on bone remodelling.

**Figure 3:**
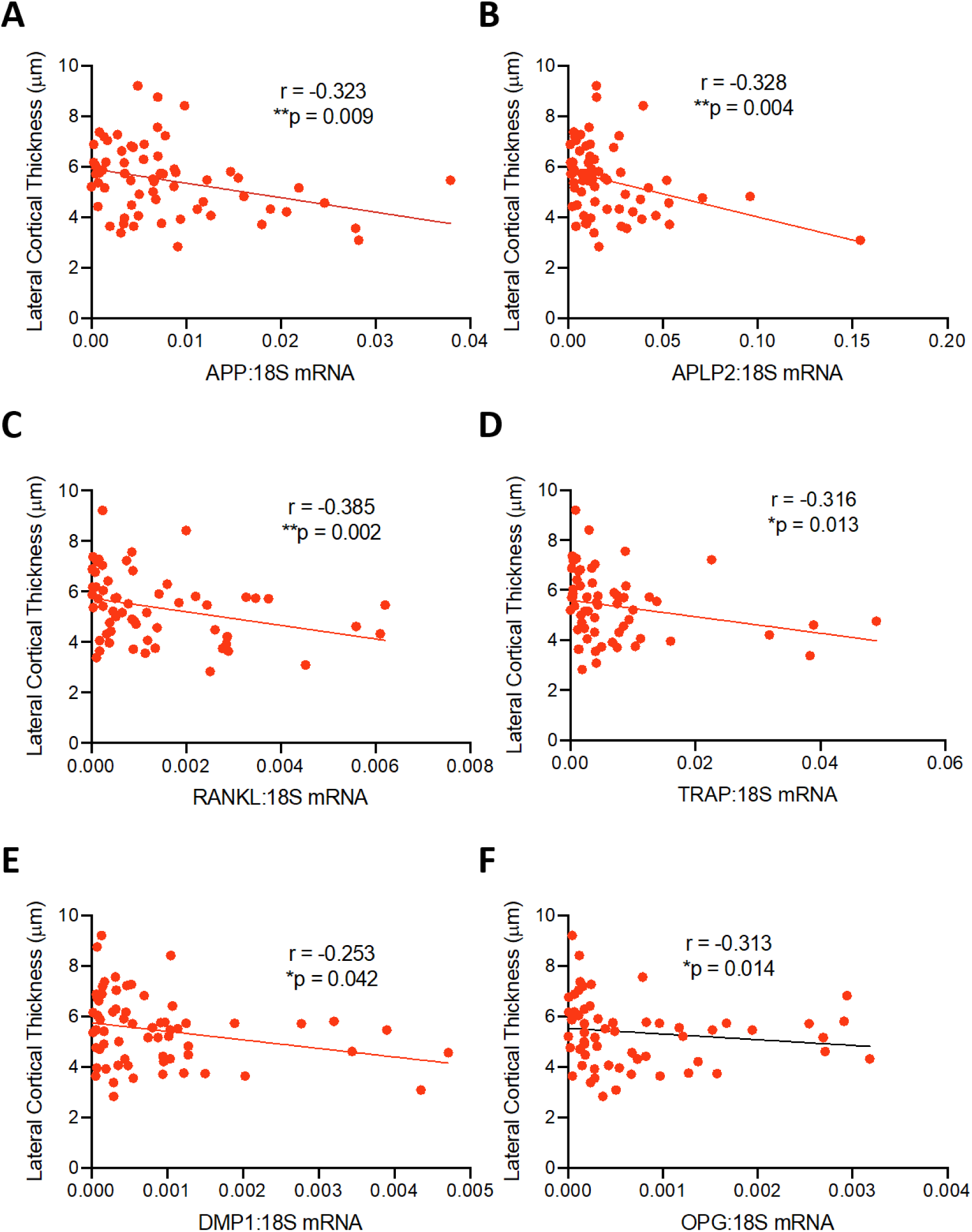
Relationships between gene expression and bone structure. Individual NOF patient radiographs were assessed for the structural parameter lateral cortical thickness, measures of which were compared with relative gene expression from the same patient, as described in Materials and Methods. Significant correlations were observed between lateral cortical thickness and A) *APP*, B) *APLP2*, C) *RANKL*, D) *TRAP*, E) *DMP1*, F) *OPG* mRNA expression. *r* values and associated values for *p* are indicated. Number of pairs = 65/parameter.

### Individual gene expression relationships between dementia and non-dementia

Next, subgroup analysis was performed based on dementia status within the NOF cohort. Subjects were dichotomised as either non-dementia or dementia, the latter based on any diagnosed form. Initial analysis demonstrated a similar pattern within each group as the entire NOF cohort, in terms relative expression of each gene. There were no significant differences between the non-dementia and dementia groups for any of the genes analysed (**Supplementary Fig. 1**).

### Gene expression and femoral structural parameter relationships between dementia and non-dementia subgroups

To determine whether relationships between neurotropic genes of interest and remodelling genes differed between non-dementia and dementia subgroups, non-parametric Spearman’s correlation analyses were performed. In the non-dementia group, there were significant correlations observed between *APP* and *APLP2* (*p* < 0.0001), *BACE1* (*p* < 0.0001), *DMP1* (*p* = 0.0009), *OPG* (*p* < 0.0001), *RANKL* (*p* < 0.0001) and *TRAP* (*p* < 0.0001). Similar correlations were observed between *APLP2* and these genes and there was also a significant correlation with *RANKL:OPG* ratio (*p* = 0.007), which was not observed with *APP. BACE1* was shown to be significantly correlated with *DMP1* (R = 0.42; *p* = 0.002), *NGF* (R = 0.54; *p* = 0.007), *OPG* (R = 0.38; *p* = 0.0078), *RANKL* (R = 0.465; *p* < 0.0001) and *TRAP* (R = 0.5; *p* = 0.0004) (**Fig. 4a**).

**Figure 4:**
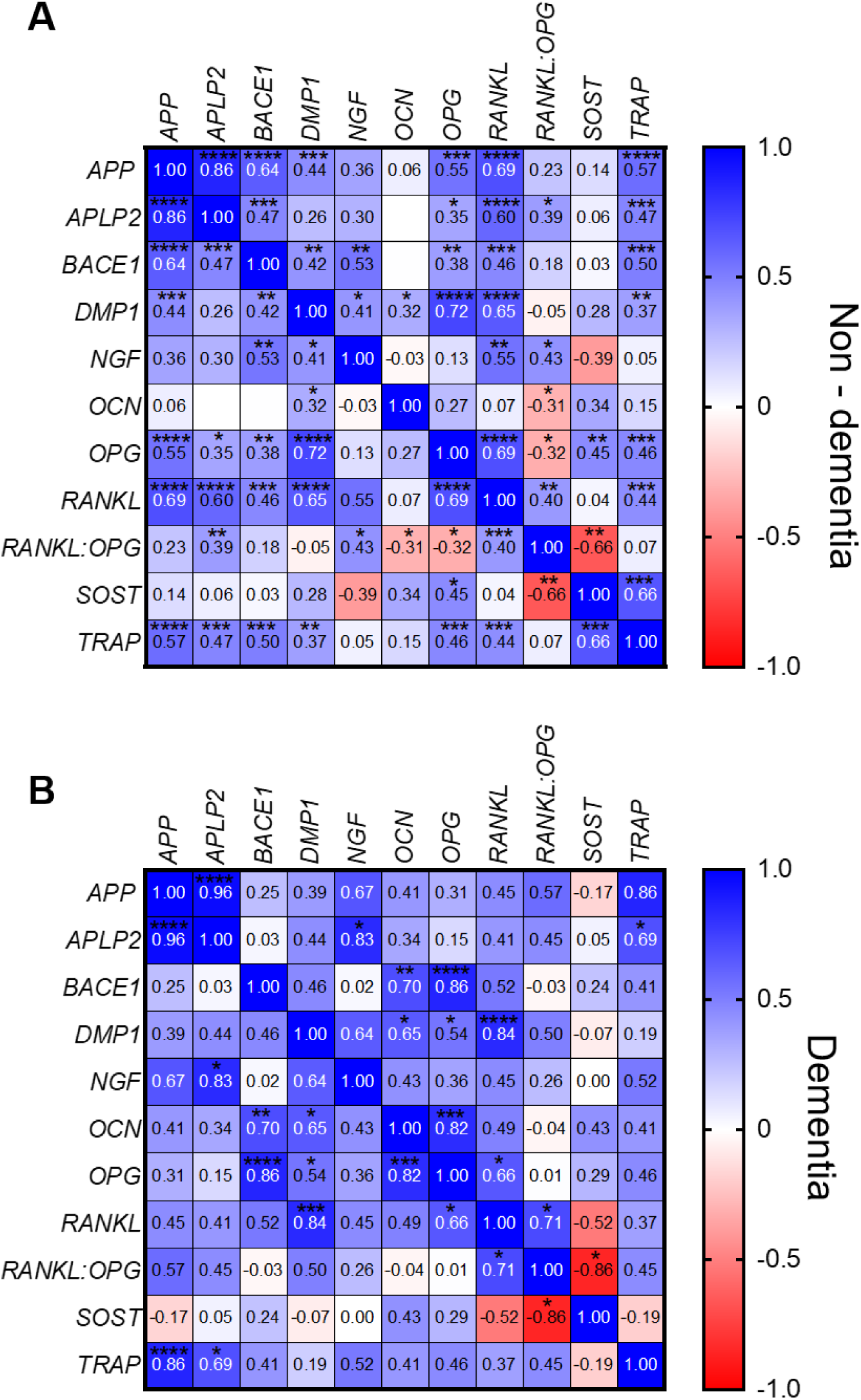
Gene expression and correlation analysis of sub-groups dichotomized on any diagnosis of dementia. Gene expression of each gene of interest was normalised to the 18S housekeeping gene. Heat map representation of the relationships between each gene of interest using the Spearman’s correlation co-efficient (*r*) for A) Non-dementia and B) Dementia sub-groups. Relationship direction and strength are indicated by the legend: 0 – 1.0 indicates positive relationship between genes and - 1.0 – 0 indicates a negative relationship between genes. The significance of *r* values is indicated by*\*p* < 0.05, \*\**p* < 0.01 and \*\*\**p* < 0.001.

In the dementia group, there were notably fewer correlations between genes of interest (**Fig. 4b**). Like in the non-dementia group, *APP* significantly correlated with *APLP2* (*p* < 0.0001), however there was no correlation with *BACE1* expression. There were weak correlations of trending significance between *APP* and *NGF* (*r* = 0.667, *p* = 0.083) and *APP* and the *RANKL:OPG* mRNA ratio (*r* = 0.573; *p* = 0.056). A strong positive correlation was, however, apparent between *APP* and *TRAP* mRNA (r = 0.857; p = 0.0004). Analysis of *APLP2* revealed a similar pattern, with a strong correlation observed with *NGF* (r = 0.83; p = 0.015) and *TRAP* (r = 0.69; p = 0.017). *BACE1* mRNA expression was significantly correlated with *OCN* (R = 0.70; *p* = 0.01) and *OPG* (*r* = 0.86; *p* < 0.0001) and the correlation between *BACE1* and *RANKL* was trending (*r* = 0.52; *p =* 0.084). Interestingly, correlations between *BACE1* and *OCN* expression were not observed in the non-dementia group. In addition, *OCN* and *OPG* were only significantly correlated in the dementia group. Finally, there was a strong negative relationship observed between *RANKL* mRNA expression and that of *SOST* in the dementia group (*r* = -0.86) (**Fig. 4b)**.

To determine whether gene expression correlated with structural femoral parameters in non-dementia and dementia groups, Spearman’s correlation testing was performed. Analysis of the non-dementia group only did not reveal any significant relationships between either *APP, APLP2 or BACE1* and structural femoral parameters (**Table 2**). There were however negative correlations observed between lateral cortical thickness and *OPG* (*r* = -0.41), *RANKL* (*r* = - 0.45) and *TRAP* (*r* = -0.35). There were also correlations observed between the structural parameters lateral cortical thickness, medial cortical thickness (*r* = 0.354; *p* = 0.021) and femoral width (*r* = 0.378; *p* = 0.014). In addition, *RANKL, RANKL:OPG* and *TRAP* mRNA expression were negatively correlated with femoral width in the non-dementia group.

**Table 2:** Correlation Analysis of Gene Expression with Structural Parameters in Non-dementia and Dementia subgroups.

| NON-DEMENTIA |  |  |  |  |  |  |  |  |
| --- | --- | --- | --- | --- | --- | --- | --- | --- |
| *Gene* | *LC (1)* |  | *MC (2)* |  | *FC (3)* |  | *PFO (4)* |  |
|  | *r* | *p* | *r* | *p* | *r* | *p* | *r* | *p* |
| *APP* | -0.246 | 0.117 | -0.252 | 0.108 | -0.090 | 0.572 | -0.247 | 0.114 |
| *APLP2* | -0.262 | 0.094 | -0.245 | 0.118 | -0.095 | 0.551 | -0.256 | 0.102 |
| *BACE1* | -0.135 | 0.396 | -0.147 | 0.353 | -0.094 | 0.554 | -0.215 | 0.172 |
| *CYP27B1* | **0.356** | **0.022** | 0.137 | 0.393 | -0.042 | 0.792 | 0.074 | 0.647 |
| *DMP1* | -0.175 | 0.267 | 0.150 | 0.343 | -0.109 | 0.492 | -0.118 | 0.458 |
| *NGF* | -0.184 | 0.425 | -0.271 | 0.234 | 0.013 | 0.955 | -0.109 | 0.638 |
| *OCN* | 0.086 | 0.589 | 0.028 | 0.862 | -0.054 | 0.734 | -0.006 | 0.970 |
| *OPG* | **-0.415** | **0.008** | -0.087 | 0.593 | 0.120 | 0.461 | -0.108 | 0.506 |
| *RANKL* | **-0.454** | **0.003** | -0.195 | 0.227 | -0.052 | 0.750 | **-0.312** | **0.050** |
| *RANKL:OPG* | -0.143 | 0.377 | -0.240 | 0.136 | -0.205 | 0.204 | **-0.316** | **0.047** |
| *SOST* | 0.192 | 0.445 | 0.158 | 0.531 | 0.238 | 0.341 | 0.269 | 0.280 |
| *TRAP* | **-0.354** | **0.025** | -0.187 | 0.249 | -0.129 | 0.428 | **-0.325** | **0.041** |
| DEMENTIA |  |  |  |  |  |  |  |  |
| *Gene* | *LC (1)* |  | *MC (2)* |  | *FC (3)* |  | *PFO (4)* |  |
|  | *r* | *p* | *r* | *p* | *r* | *p* | *r* | *p* |
| *APP* | -0.624 | 0.060 | -0.224 | 0.537 | 0.261 | 0.470 | -0.212 | 0.560 |
| *APLP2* | -0.479 | 0.166 | -0.286 | 0.420 | 0.418 | 0.233 | 0.091 | 0.811 |
| *BACE1* | -0.264 | 0.435 | -0.087 | 0.802 | 0.045 | 0.903 | -0.309 | 0.356 |
| *CYP27B1* | -0.236 | 0.513 | -0.297 | 0.407 | 0.200 | 0.584 | 0.200 | 0.584 |
| *DMP1* | -0.418 | 0.233 | -0.559 | 0.098 | 0.200 | 0.584 | -0.552 | 0.105 |
| *NGF* | -0.500 | 0.267 | -0.107 | 0.840 | 0.214 | 0.662 | 0.179 | 0.713 |
| *OCN* | -0.345 | 0.299 | -0.114 | 0.739 | 0.064 | 0.860 | -0.309 | 0.356 |
| *OPG* | -0.073 | 0.838 | 0.091 | 0.790 | -0.155 | 0.654 | -0.327 | 0.327 |
| *RANKL* | -0.273 | 0.448 | -0.219 | 0.541 | 0.055 | 0.892 | -0.503 | 0.144 |
| *RANKL:OPG* | -0.176 | 0.632 | 0.164 | 0.649 | -0.018 | 0.973 | -0.261 | 0.470 |
| *SOST* | -0.286 | 0.556 | -0.536 | 0.236 | 0.429 | 0.354 | 0.286 | 0.556 |
| *TRAP* | -0.391 | 0.237 | 0.178 | 0.599 | 0.200 | 0.557 | 0.173 | 0.615 |

Analysis of the dementia group revealed a trending negative correlation between *APP* and lateral cortical thickness (*r* = -0.591; *p* = 0.061). Analysis of the femoral regions revealed strong correlations between lateral cortical thickness and medial cortical thickness (*r* = 0.597; *p* = 0.057) and a negative correlation with medullary width (*r* = -0.782; *p* = 0.006) (**Table 2**).

### Relationships between genes of interest and serum Vitamin D or creatinine

Analyses of all genes against serum Vitamin D and creatinine levels were performed to determine if there were differences between dementia and non-dementia groups. There were no significant correlations between *APP, APLP2 or BACE1* and Vitamin D or creatinine. In both the non-dementia and dementia groups however, there were correlations between *OPG* mRNA expression and serum creatinine levels (**Supplementary Fig. 2**).

## Discussion

Osteoporosis leading to fragility fracture of the hip and dementia are known to afflict a similar demographic. These diseases not only contribute to high rates of morbidity and mortality in the aged but also place a huge economic burden on the health care system. Patients undergoing arthroplasty for the management of their fracture are a diverse cohort of a wide age range, who suffer from a myriad of comorbidities, which makes them a difficult group to assess at the molecular level. In keeping with this being a ‘silent disease’ [5], only 18% of patients in this study had a pre-existing diagnosis of osteoporosis. This is one of the first cohort studies combining bone genetics, bone structure, systemic measures and co-morbidity information in the NOF population in Australia in an attempt to uncover links between these pathologies.

Whole cohort analysis revealed for the first time strong coordinated expression of the CNS related genes *APP, APLP2* and *BACE1* in adult human bone tissue. Intriguingly, the expression of these genes was closely related to known markers of bone turnover, *RANKL* and *TRAP*, and osteoblast activity, *OPG* and *DMP1*. The strongest correlations were observed between *APP, APLP2, BACE1* and *RANKL* and *TRAP* not only in the whole cohort analysis, but also within the dementia subgroup. These strong associations are suggestive of a linked functionality between genes involved in CNS homeostasis and pathology in bone remodelling. In the context of NOF fracture, the relationships were still existent in the non-dementia group as well as the dementia group, indicating that there may be an acceleration of bone loss in this subgroup, however this warrants further analysis of a larger cohort powered to determine differences between various classifications of cognitive impairment. Despite many of the relationships seen in both the non-dementia and dementia groups being consistent, there were unexpected relationships between *BACE1, OCN* and *OPG* observed solely in the dementia group. These correlations unique to dementia bone specimens may indicate possible alternate mechanisms of bone loss occurring. These observations may indicate a role for genes involved in the homeostasis of the CNS in processes such as bone resorption through either direct or indirect mechanisms. However, as these are associations, it is not possible to determine the driver of each relationship observed.

The strong relationship evident between serum creatinine and *OPG* mRNA expression in the whole NOF cohort was an unexpected finding. Previous relationships between serum OPG and creatinine have been noted in the context of severe artery calcification [43], where increased serum OPG was found to be a potential marker for all-cause mortality in patients with chronic kidney disease. This relationship requires further elucidation in the context of fracture as it may be an indicator of other comorbidities.

Analysis of the femoral cortical bone parameters is a useful tool to assess relationships between bone structure and gene expression. The cortical thickness index has been tested as a surrogate for BMD in hip fracture patients [44-46]. In this study, the structural parameter most closely related to the largest number of functional genes was lateral cortical thickness. A similar pattern was observed within the whole cohort with inverse relationships between the expression of all of *RANKL, TRAP, OPG* and *DMP1* and lateral cortical thickness, suggesting that all of these genes known to be related to bone remodelling are linked to increased bone loss or reduced bone formation at this particular site. In particular, *TRAP* expression was strongly correlated with a number of different genes and parameters, and can be a marker of not only osteoclastic resorption but also osteocytic osteolysis [24]. This highlights the need to investigate further the role of osteocytes in age-related bone loss and potential accelerated bone loss in dementia. Interestingly, the expression of *APP* and *APLP2* were also correlated with this structural parameter, implicating these neurotropic genes in the control of bone structure. In the subgroup analysis, despite the small number of participants in the dementia group, these trends were still observed. The inverse relationships between *APP* and *APLP2* and cortical thickness suggests positive roles for these genes in net bone loss. Future studies are warranted to determine if these genes contribute to cortical thinning or cortical porosity, characteristic of osteoporosis and fracture risk [47].

As this was a study solely of a population undergoing emergency total hip replacement surgery for NOF, we cannot determine whether the relationships observed are specific to NOF or are generalizable to the population as a whole. However, in a time where average life expectancies and the ageing population are increasing, it is imperative to understand the genetic changes that are occurring in the bone in these later stages of life. In particular, links between cognitive decline and systemic manifestations of ageing and frailty, such as osteoporosis, need to be better understood.

## Declarations

The study was approved by the Human Research Ethics Committee of the RAH (Protocol no 130114; HREC/13/RAH/33) and all participants provided written, informed consent. All authors contributed to this study and approve the final version. GJA was supported by a National Health and Medical Research Council of Australia (NHMRC) Senior Research Fellowship. CJMS was supported by an Australian Postgraduate Award scholarship.

## Acknowledgements

The authors thank the nursing and surgical staff of the Orthopaedic and Trauma Service of the Royal Adelaide Hospital for help in obtaining patient samples.

---

**Supplementary Figure 1:** Gene expression analysis for non-dementia and dementia groups. Real-time RT-PCR analysis of genes of interest normalised to the 18S housekeeping gene for non-dementia group (*n* = 53) versus the dementia group (*n* = 13). There were no statistical differences between groups for any gene (Student’s t-test). Data are depicted as means only.

**Supplementary Figure 2:** Correlations between *OPG* mRNA expression and serum creatinine in non-dementia and dementia groups. A. *OPG* mRNA expression was moderately correlated with serum creatinine in the non-dementia group (n = 48). B. *OPG* and serum creatinine were positively correlated in the dementia group (n = 13).

