## Supplementary Figure 1 for "Relationships between the Bone Expression of Alzheimer Disease-Related Genes, Bone Remodelling Genes and Cortical Bone Structure in Neck of Femur Fracture"

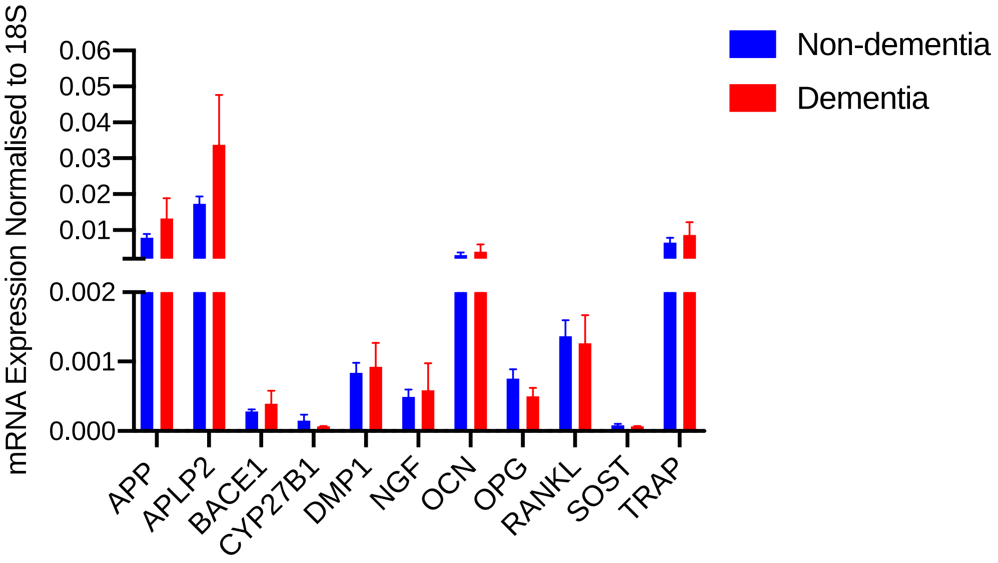


**Supplementary Figure 1:** Gene expression analysis for non-dementia and dementia groups. Real-time RT-PCR analysis of genes of interest normalised to the 18S housekeeping gene for non-dementia group ($n$ = 53) versus the dementia group ($n$ = 13). There were no statistical differences between groups for any gene (Student's t-test). Data are depicted as means only.
