## Supplementary Figure 2 for "Relationships between the Bone Expression of Alzheimer Disease-Related Genes, Bone Remodelling Genes and Cortical Bone Structure in Neck of Femur Fracture"

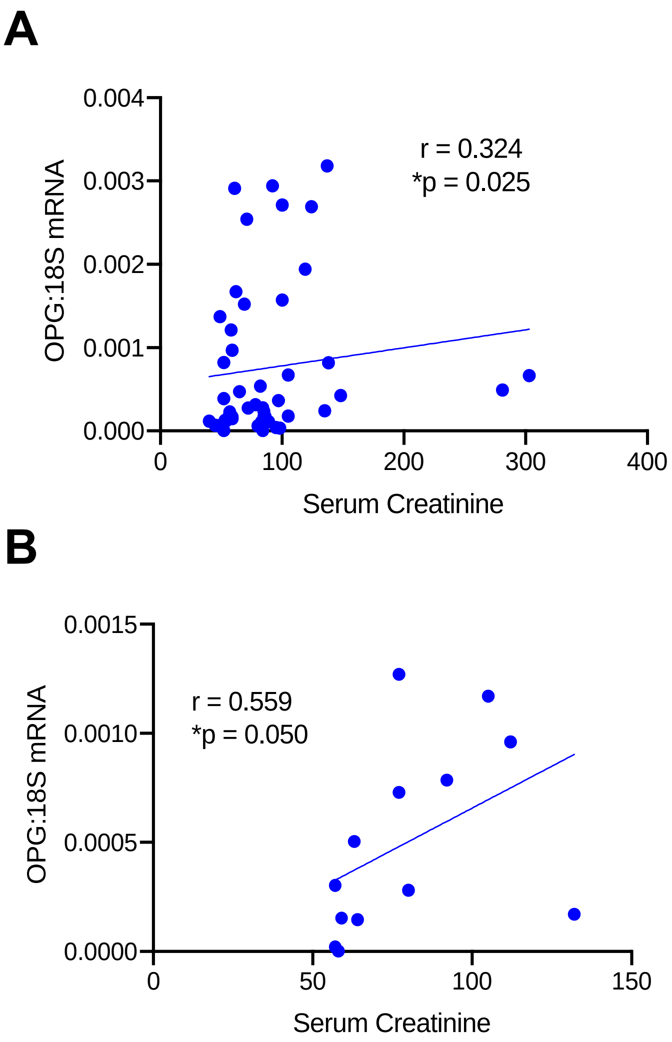


**Supplementary Figure 2:** Correlations between *OPG* mRNA expression and serum creatinine in non-dementia and dementia groups. A. *OPG* mRNA expression was moderately correlated with serum creatinine in the non-dementia group (n = 48). B. *OPG* and serum creatinine were positively correlated in the dementia group (n = 13).
