## Supplementary Tables 1-3 for "Relationships between the Bone Expression of Alzheimer Disease-Related Genes, Bone Remodelling Genes and Cortical Bone Structure in Neck of Femur Fracture"

**Supplementary Table 1. Forward and reverse primer sequences for real-time RT-PCR**

| Gene | Direction | Primer Sequences (5´ – 3´) |
|---|---|---|
| ***18S*** | F | GGAATTCCCGAGTAAGTGCG |
| | R | GCCTCACTAAACCATCCAA |
| ***APP*** | F | ATCCTGCAGTATTGCCAAGAAG |
| | R | CACAAAGTGGGGATGGGTC |
| ***APLP2*** | F | GCCCAGATGAAATCCCAGGT |
| | R | ATATCTGCACGCTGCTCCTG |
| ***BACE1*** | F | GCAGGGCTACTACGTGGAGA |
| | R | GTATCCACCAGGATGTTGAGC |
| ***CYP27B1*** | F | TGGCCCAGATCCTAACACATTT |
| | R | GTCCGGGTCTTGGGTCTAACT |
| ***DMP1*** | F | GATCAGCATCCTGCTCATGTT |
| | R | AGCCAAATGACCCTTCCATTC |
| ***NGF*** | F | CACACTGAGGTGCATAGCGT |
| | R | TGATGACCGCTTGCTCCTGT |
| ***OCN*** | F | TGAGAGCCCTCACACTCCTC |
| | R | ACCTTTGCTGGACTCTGCAC |
| ***OPG*** | F | GCTCACAAGAACAGACTTTCCAG |
| | R | CTGTTTTCACAGAGGTCAATATCTT |
| ***RANKL*** | F | CCAAGATCTCCAACATGACT |
| | R | TACACCATTAGTTGAAGATACT |
| ***SOST*** | F | ACCGGAGCTGGAGAACAACA |
| | R | GCTGTACTCGGACACGTCTT |
| ***TRAP*** | F | GTGCAGACTTCATCCTGTCTCTA |
| | R | AATACGTCCTCAAAGGTCTCC |

**Supplementary Table 2: Cohort demographics and pre-fracture comorbidities.**

| *Cohort Demographics* | *n* | *Prevalence (%)* | *Age (mean ± SD)* |
|---|---|---|---|
| ***Female*** | 53 | 80.30 | 82.4 (9.15) |
| ***Male*** | 13 | 19.70 | 80.2 (9.28) |
| *Whole Cohort* | 66 | 100.0 | 81.9 (9.15) |

| *Comorbidity* | ***n*** | ***Prevalence (%)*** | ***Comorbidity*** | ***n*** | ***Prevalence (%)*** |
|---|---|---|---|---|---|
| ***Hypertension*** | 43 | 65.15 | *Anaemia* | 4 | 6.06 |
| ***Hypercholesterolemia*** | 20 | 30.30 | *Alcohol Abuse* | 3 | 4.55 |
| ***Osteoarthritis*** | 17 | 25.76 | *Aortic Stenosis* | 3 | 4.55 |
| ***Type 2 Diabetes Mellitus*** | 17 | 25.76 | *Asthma* | 3 | 4.55 |
| ***Atrial Fibrillation*** | 16 | 24.24 | *Cardiac Stent* | 3 | 4.55 |
| ***Gout*** | 16 | 24.24 | *Glaucoma* | 3 | 4.55 |
| *Dementia* | 13 | 19.69 | *Paroxysmal AF* | 3 | 4.55 |
| *Osteoporosis* | 12 | 18.18 | *Recurrent Falls* | 3 | 4.55 |
| ***GORD*** | 11 | 16.67 | *Smoker* | 3 | 4.55 |
| ***Depression*** | 8 | 12.12 | *Total Knee Replacement* | 3 | 4.55 |
| ***Hypothyroidism*** | 8 | 12.12 | *Peripheral Vascular Disease* | 3 | 4.55 |
| ***Chronic Obstructive Pulmonary Disease*** | 7 | 10.61 | *Angina* | 2 | 3.03 |
| ***Recurrent UTI*** | 7 | 10.61 | *Aortic Valve Replacement* | 2 | 3.03 |
| ***Anxiety*** | 5 | 7.58 | *Breast Cancer* | 2 | 3.03 |
| ***Macular Degeneration*** | 5 | 7.58 | *Hemi arthroplasty* | 2 | 3.03 |
| ***Myocardial Infarction*** | 5 | 7.58 | *Hiatus Hernia* | 2 | 3.03 |
| ***Hearing Impairment*** | 4 | 6.06 | *Ovarian Cancer* | 2 | 3.03 |
| ***Cholecystectomy*** | 4 | 6.06 | *Parkinson's Disease* | 2 | 3.03 |
| ***Chronic Kidney Disease*** | 4 | 6.06 | *Pulmonary Embolism* | 2 | 3.03 |
| ***Stroke*** | 4 | 6.06 | *Thyroid Cancer* | 2 | 3.03 |
| ***Diverticulosis*** | 4 | 6.06 | *Urinary Incontinence* | 2 | 3.03 |
| ***Hysterectomy*** | 4 | 6.06 | | | |

**Supplementary Table 3: Correlation analysis between serum Vitamin D and serum creatinine and genes of interest in the whole NOF cohort.**

| **Gene** | **Vitamin D (*R*)** | **Creatinine (*R*)** |
|---|---|---|
| ***APP*** | -0.031 | 0.025 |
| ***APLP2*** | 0.017 | -0.043 |
| ***BACE1*** | 0.133 | 0.170 |
| ***CYP27B1*** | 0.109 | -0.038 |
| ***DMP1*** | 0.096 | 0.064 |
| ***NGF*** | -0.102 | 0.179 |
| ***OCN*** | 0.234 | -0.001 |
| ***OPG*** | 0.039 | 0.346** |
| ***RANKL*** | 0.009 | 0.150 |
| ***RANKL:OPG*** | -0.025 | -0.154 |
| ***SOST*** | 0.260 | -0.231 |
| ***TRAP*** | 0.113 | 0.014 |

Significance is indicated by **$p < 0.01$
